# Increased stromal densities of B cells, CD103+ cells, and CD163+ M2-like macrophages associate with poor clinical outcomes in BCG treated non-muscle invasive bladder cancer

**DOI:** 10.1101/2023.10.19.562817

**Authors:** Benjamin Ravenscroft, Priyanka Yolmo, Stephen Chenard, Sadaf Rahimi, Keiran Pace, Kartik Sachdeva, Tamara Jamaspishvilli, Hamid Ghaedi, Andrew Garven, Kathrin Tyryshkin, David M. Berman, Chelsea Jackson, D. Siemens Robert, Madhuri Koti

## Abstract

Non-muscle invasive bladder cancer (NMIBC) constitutes a significant clinical challenge, with over 50% of patients experiencing poor clinical outcomes in the form of early recurrence or progression following treatment with Bacillus Calmette-Guerin (BCG) immunotherapy. The pre-treatment tumor immune microenvironment (TIME) is an established determinant of response to BCG. This study explores the spatial profiles of CD79a+ B cells, CD163+ M2-like macrophages, proliferating and tissue-resident phenotypes of T cells, along with PD-1/PD-L1 checkpoint expression in pre-BCG treatment tumors of 173 patients (139 males, 34 females). Multiplex immunofluorescence staining of a tumor tissue microarray, revealed elevated infiltration of CD79a+ B cells, CD163+ M2-like macrophages, CD103+ cells, and CD8+ T cells at the tumor invasive margins. Increased epithelial PD-L1 immune-checkpoint expression in tumors was observed in female and male patients who exhibited significantly shorter recurrence-free survival (RFS). Importantly, high CD79a+ B cell density in BCG-treated females in both stromal and epithelial compartments exhibited significantly shorter RFS and progression-free survival compared to males. Stromal CD79a+ B cell density was positively correlated with M2-like macrophages, CD8+ T cells, CD103+ cells and PD-1 expressing cells. CD79a+ B cells, CD103+ cells, and M2-like macrophage density were associated with higher grade and enriched in basal subtype tumor. This study highlights the significance of an understudied role of B cells and their cellular neighborhoods in the pre-treatment TIME and BCG-therapy response. Overall, findings from this study underscore the importance of considering sex-related immunobiological differences in the stromal compartments of bladder tumors towards the development of optimal therapeutic targeting strategies.

## Introduction

Approximately 75% of newly diagnosed bladder cancer patients present with NMIBC. While the incidence of NMIBC is four times higher in males, female patients with NMIBC often present with a more advanced stage and display earlier recurrence [1–4]. Patient age, tobacco smoking, and occupational exposure to chemicals are some of the major risk factors associated with NMIBC [5–7]. Bacillus Calmette-Guérin (BCG), a live attenuated strain of *Mycobacterium bovis*, is the adjuvant treatment for patients with intermediate or high-risk NMIBC [8, 9]. Intravesical BCG therapy entails 6 weekly instillations after transurethral resection of bladder tumor (TURBT), followed by a maintenance schedule of 3 weekly instillations every 3-6 months for 1-3 years depending on the risk stratification as defined by the American and European Urological Association [9]. Despite its proven efficacy, over 50% of the patients receiving BCG immunotherapy experience recurrence or become unresponsive to treatment [5, 6]. Second line treatments such as programmed cell death protein-1 receptor/ligand (PD-1/PD-L1) immune checkpoint axis inhibitors and replication deficient adeno viral gene-delivery vectors (Adstiladrin) are under trials for BCG unresponsive/recurrent disease [10–12]. While some have demonstrated modest benefits, these newer treatment approaches do not increase primary response to BCG and are not indicated in the BCG naïve setting [13–15].

The pre-treatment TIME is a major determinant of response to immunotherapy across various cancers [11, 16, 17]. Recent studies have established that the pre-treatment TIME plays an important role in determining BCG therapy response in patients with NMIBC [18–21]. This is especially important when considering BCG therapy, which involves a live bacterial preparation and its efficacy is dependent on multiple factors including the dose, viability of bacteria at the time of instillation, immune physiological status of the host, bacterial contact with the urothelium and uptake by cells in the bladder mucosal immune environment. Based on the infiltration profiles of CD8+ cytotoxic T cells, FoxP3+ T_reg_ cells, and the expression patterns of PD-1/PD-L1+ immune checkpoints, both in pre-BCG treatment and recurrent tumors, a potential role of adaptive immune resistance has also been suggested [19]. Additionally, elevated T cell exhaustion has been reported as a major factor underlying poor response to BCG [20]. In our previous report, conducted on the whole transcriptomic profiles of the UROMOL cohort (n=460) and spatial immunophenotyping of tumors from a large cohort assembled at the Kingston Health Sciences Center (KHSC, n=332), we showed increased infiltration of B cells, M2-like macrophages, and higher expression of PD-L1 in the pre-treatment TIME of both female and male patients experiencing early recurrence, irrespective of the treatment type [18]. Notably, female patients with high-grade disease in both cohorts exhibited significantly shorter progression free survival (PFS) and had higher infiltration of B cells and M2-like macrophages in their pre-treatment tumors compared to their male counterparts. In the current study, we characterized the association between immune cell spatial profiles in the BCG-naïve TURBT specimens of female and male patients exhibiting differential response to BCG immunotherapy, from our previously reported KHSC cohort [18]. Given the established differences in sex-associated B cell immunobiology and the need for improved patient stratification for increased benefits from BCG immunotherapy, we further evaluated patient sex-associated pre-BCG tumor immune profiles and clinical outcomes.

## METHODS

### Ethics statement

This study was approved by the Queen’s University Health Sciences and Affiliated Teaching Hospitals Research Ethics Board.

### *In silico* analysis of *CD79A, CD103* and *CD163* gene expression profiles in transcriptomic profiles of tumors from BCG naïve high-risk patients

Based on our previous findings [18] on the associations between tumor infiltrating CD79a+ B cells and M2-like macrophage densities and overall poor clinical outcomes in patients with NMIBC, irrespective of the treatment type, we first evaluated the publicly available normalized bulk RNA-sequencing based tumor whole transcriptome profiles of 283 patients with NMIBC that was recently generated by Jong *et al.,* [21]. The patients in this cohort were stratified into different quartiles according to their *CD79a* and *CD163* gene expression levels. Patient clinical demographics and outcome data was extracted from the supplementary information provided by de Jong *et al*., [21] These clinical attributes were then combined with the gene expression profiles of the primary tumors. Subsequently, an analysis was conducted using the time to disease progression and the progression status in R version 4.2.0. To visualize the relationship between target *gene* expression and survival, Kaplan-Meier survival curves were generated by employing a log-rank statistical analysis method and utilizing the survival and survminer packages in R.

### KHSC-BCG Patient Cohort and Clinical Data

Given their critical roles in mucosal immunobiology we then asked whether the spatial profiles of CD79a+ B cells, CD163+ M2-like macrophages and other immune cells of interest (see below) associate with variable response to BCG in female and male patients with NMIBC. Of the 332 patients reported in our previous study [18], a total of 173 patients (KHSC-BCG:139 males and 34 females) who received at least one induction round of BCG, defined as at least 5/6 instillations, and had no BCG treatment prior to entering the cohort, were selected for inclusion in this study (**Table 1**). Adequate BCG treatment was defined as an initial induction round of at least 5/6 instillations, followed by a maintenance round of at least 2/3 instillations [9]. Patients were assessed for BCG therapy response and classified as either BCG refractory, BCG responders, or an in-between state of therapy response. The BCG refractory state was defined according to the recent consensus guidelines put forth by the European Association of Urology [6], and BCG responders were patients without any high-grade recurrence within the two years, following their first adequate BCG induction (**Table 1**)[8]. Due to our cohort size limitations and the differences in clinical treatment versus ideal clinical trial treatment pathways, we used high-grade recurrence as an endpoint for time-to-event analysis. The initiation point for time-to-event analyses is the diagnosis of TURBT immediately prior to the initiation of BCG treatment. Progression events were defined as recurrence to stage T2 or higher.

**Table 1:** Categories of BCG therapy response.

|  |
| --- |
| **BCG-refractory tumour (as per standard definitions)[8]** |
| 1. If T1 G3/HG tumour is present at 3 mo. |
| 2. If TaG3/HG tumour is present after 3 months and/or at 6 mo., after either re-induction or first course of maintenance. |
| 3. If CIS (without concomitant papillary tumour) is present at 3 mo. and persists at 6 mo. after either reinduction or a first course of maintenance. For patients with CIS present at 3 mo., an additional BCG course can achieve a complete response in >50% of cases. |
| 4. If HG tumour appears during BCG maintenance therapy. |
| **BCG responsive** |
| 1. No HG tumour recurrence within 2 years following initial adequate BCG induction. |

Baseline clinical data and NMIBC tissue were collected for eligible patients (**Table 2**). All the tumor tissue specimens included in the study were index TURBT specimens. The median follow-up from surgical resection (TURBT) was 54.3 months (95% CI 50.2 to 59.5 months) for the cohort. 124 (71.7%) of the cohort were BCG responders and 37 (21.4%) were BCG refractory. The entire cohort had a 24-month high-grade recurrence free survival (RFS) rate of 70% (95% CI 63.1% to 77.5%). The median time to HG recurrence was 7.6 months (95% CI 6.3 to 11.8 months), and the median follow-up time for BCG responders was 53.4 months (95% CI 47.8 to 59.5 months).

**Table 2:** Patient cohort clinical characteristics.

|  | BCG<br>Responsive<br>(N=124) | BCG Refractory<br>(N=37) | BCG relapsing<br>(N=12) | Overall<br>(N=173) |
| --- | --- | --- | --- | --- |
| **Age** |  |  |  |  |
| Mean | 71.5 | 68.3 | 72.3 | 70.8 |
| Median | 71.5 | 69.0 | 73.0 | 71.0 |
| Range | 35-93 | 45-85 | 62-82 | 35-93 |
| **Sex** |  |  |  |  |
| Male | 102 (82.3%) | 27 (73.0%) | 10 (83.3%) | 139 (80.3%) |
| Female | 22 (17.7%) | 10 (27.0%) | 2 (16.7%) | 34 (19.7%) |
| **AUA Risk Score** |  |  |  |  |
| High | 74 (59.7%) | 36 (97.3%) | 12 (100.0%) | 122 (70.5%) |
| Intermediate | 41 (33.1%) | 1 (2.7%) | 0 (0%) | 42 (24.3%) |
| Low | 9 (7.3%) | 0 (0%) | 0 (0%) | 9 (5.2%) |
| **Diagnosis TURBT**<br>**Stage and Grade** |  |  |  |  |
| CIS | 11 (8.9%) | 1 (2.7%) | 1 (8.3%) | 13 (7.5%) |
| PUNLMP | 1 (0.8%) | 0 (0%) | 0 (0%) | 1 (0.6%) |
| T1HG | 32 (25.8%) | 15 (40.5%) | 4 (33.3%) | 51 (29.5%) |
| TaHG | 51 (41.1%) | 17 (45.9%) | 5 (41.7%) | 73 (42.2%) |
| TaLG | 29 (23.4%) | 2 (5.4%) | 2 (16.7%) | 33 (19.1%) |
| Unknown | 0 (0%) | 2 (5.4%) | 0 (0%) | 2 (1.2%) |
| **Time to Follow-up** |  |  |  |  |
| Median time to<br>follow-up (months) | 53.4 | 57.9 | 65.4 | 54.3 |
| **BCG Adequacy** |  |  |  |  |
| Adequate BCG | 47 (37.9%) | 23 (62.2%) | 11 (91.7%) | 81 (46.8%) |
| Induction only | 77 (62.1%) | 14 (37.8%) | 1 (8.3%) | 92 (53.2%) |
| **Subtype** |  |  |  |  |
| Genomically<br>Unstable | 13 (10.3%) | 3 (7.7%) | 1 (8.3%) | 17 (9.6%) |
| URO-KRT5+ | 11 (8.7%) | 2 (5.1%) | 2 (16.7%) | 15 (8.5%) |
| Urothelial-like | 9 (7.1%) | 7 (17.9%) | 1 (8.3%) | 17 (9.6%) |
| Basal | 1 (0.8%) | 1 (2.6%) | 0 (0.0%) | 2 (1.1%) |
| Not Subtyped | 92 (73.0%) | 26 (66.7%) | 8 (66.7%) | 126 |
| **Subtype Grade** |  |  |  |  |
| 1 | 27 (21.4%) | 7 (17.9%) | 2 (16.7%) | 36 (20.3%) |
| 2 | 7 (5.6%) | 6 (15.4%) | 2 (16.7%) | 15 (8.5%) |
| Not Subtyped | 92 (73.0%) | 26 (66.7%) | 8 (66.7%) | 126 |

### Multiplex Immunofluorescence Staining

Three panels of antibodies were used to stain the tissue microarrays (TMAs; **Supplementary data, Figure S1**). The first panel consisted of antibodies identifying FoxP3+, CD8+, CD3+, and Ki67+ cells. The second panel PD-1+, PD-L1+, and CK5+ cells and the third panel included antibodies identifying GATA3+, CD103+, CD163+, and CD79a+ cells. The following cell types were evaluated for expression in both the epithelium and stroma: proliferating and non-proliferating cytotoxic T cells (CD3+CD8+Ki67+/-), cytotoxic regulatory and non-regulatory T cells (CD3+FoxP3+/-), immune checkpoints (PD-1+, PD-L1+, PD-L1+ expressed on CK5+ cancer cells and immune cells), CD103+ cells, CD163+ M2-like macrophages and CD79a+ B cells. Staining and scanning of the TMA was performed using multiplex immunofluorescence (mIF) staining at the Molecular and Cellular Immunology Core (MCIC) facility, BC Cancer Agency as described earlier [18]. The antibodies utilized were supplied by Biocare Medical (Pacheco, CA, USA) and distributed by Inter Medico (Markham, ON, CAN).

### Automated scoring and manual validation of immune cell density and distribution in mIF stained tumors

The stained TMA sections were scanned using the Vectra 3.0.5 multispectral imaging system (PerkinElmer, MA, USA) and subsequently analyzed in InForm 2.4.0 (PerkinElmer, MA, USA). The scanned TMAs underwent tissue segmentation into epithelium and stroma utilizing InForm’s tissue segmentation algorithm, specifically examining nuclear DAPI. Phenotyping was performed by training three independent algorithms on ten randomly selected cores from the cohort. The software identified positive pixels for all selected immune markers using each of the three independent algorithms. The average was taken across the three algorithms for all three sets of immune markers in each core. Cores containing ≤ 25% of the tissue were excluded from subsequent analyses. Standard deviation between the three algorithms was calculated for each of the immune markers within both the epithelial and stromal compartments. Outliers were identified and cross-referenced with the composite image of the corresponding TMA core to verify the discrepancy before excluding the outlying data points from further analysis. Any quantifications that included misidentification of histological artifacts as immune cells by any of the three algorithms were also excluded from analysis. Algorithms that consistently over-called or under-called the immune marker quantification were excluded from analysis. Further visual validation of the automated scoring was performed for randomly selected TMA cores.

### Statistical Analysis

All analyses were conducted using either R version 4.1.2 or Python 3.9.12. Analysis of cell population composition was performed using unpaired non-parametric Mann-Whitney tests. The corrplot package was utilized to calculate and visualize the Spearman rank correlations of various cell type, due to its non-parametric nature making it better suited for data lacking normality. The Kaplan-Meier method and log-rank test were performed to determine the association between time to high-grade recurrence and dichotomized infiltration of immune cell phenotypes.

Differences in infiltration levels of immune cells between BCG refractory patients and BCG responders were tested and presented with violin plots. Prior to statistical analysis, the normality condition was checked for the density of each phenotype across the whole cohort using the Shapiro-Wilk test. Failing to satisfy this condition, the Wilcox-rank sum test was used to test for differences in immune cell infiltration in BCG responders and non-responders as it does not assume normality.

To dichotomize infiltration of specific immune cell phenotypes, a combination of graphical diagnostic plot methods as presented by Williams *et al.* and bootstrapping were used (<u>Microsoft Word - Technical report 79 cover mandrekar 6-06.doc (mayo.edu)</u>. Unique densities of each immune phenotype were used to create indicator variables that could fit Cox models, from which log-likelihood values could be extracted. These values were plotted, and LOESS curve fitting was performed using the stats package. The discrete infiltration values closest to the five most visually unique peaks were selected. Bootstrapping was performed using a univariable cox model that compared the high/low infiltration groups for each of the six possible cut point values previously identified [18]. The validated cut point was the dichotomization value with the greatest frequency of maximum log-likelihood after bootstrapping. Valid cut points were used to create dichotomized Kaplan-Meier curves comparing RFS and PFS between high and low infiltration groups for various immune cell phenotypes. The dichotomized Kaplan-Meier curves were typically truncated prior to the number at risk in any group dropping below 5.

The distribution of clinically prognostic variables previously evaluated [18], including subtype, grade, stage, AUA risk score, sex, and age, was rigorously examined for each dichotomized immune cell phenotype. Statistical analysis involved employing Pearson’s χ² test or Fisher’s Exact test in cases where all assumptions were not met. P-values were calculated both without adjustment for multiple testing and with adjustment for multiple testing using the Bonferroni method.

The clinical impact of immune cell and immune checkpoint populations were analyzed using a univariable Cox proportional hazards (CPH) model. The proportional hazards (PH) assumption was verified using the Schoenfeld test, computed using the survival package. A nonsignificant (p > 0.05) test statistic indicates that the PH assumption is met. Multivariable models were created for each immune cell phenotype with age, sex, AUA risk score, and BCG treatment amount, dichotomized into induction only and adequate BCG treatment groups, as covariates. P-values were calculated with and without adjusting for multiple testing using the Bonferroni method for correction of multiple testing. Results are reported graphically in a forest plot and hazard ratios (HR) for immune cell density variables are scaled by the one standard deviation of the given immune cell population when graphically reported. Scaling the hazard ratios by each immune cell population’s standard deviation in the cohort was performed to enable clearer understanding of relative association between each cell type and RFS/PFS, as well as comparison of each different cell type’s effect size. Some cell types have a very low SD in the cohort, such as stromal CK5+PD-L1+ (SD = 5.06), while others, such as stromal CD79a+ B cells (SD = 137.13), have a very large standard deviation. Subsequently, reporting HR scaled by SD (hereafter denoted HR_SD_) shows effect size relative to expected magnitude of change in cell infiltration for a given cell type [22].

## RESULTS

### Increased expression of CD79A and CD163 genes in pre-treatment tumors is associated with shorter progression free survival following treatment with BCG

We first determined the association between expression levels of *CD79A* and *CD163* transcripts and clinical outcomes using the publicly available and recently published large independent cohort of BCG treated patients [21]. In this cohort of 283 patients, 20% of the patients were female and the entire cohort was BCG-naive high-risk NMIBC with 34% of the patients exhibiting progression after treatment with BCG. Kaplan-Meier survival analysis revealed that high expression of *CD79A and CD163 are* associated with significantly shorter PFS following BCG therapy (**Supplementary data, Figure S2**). In this cohort, this association was observed in both male and female patients (not shown).

### High CD79a+ B cell infiltration in stromal and epithelial regions is associated with early recurrence and progression following BCG therapy

The patient clinical demographics of KHSC-BCG cohort are presented in **Table 2**. Of the 173 patients in the KHSC-BCG cohort, 72% patients had a diagnosis of high-grade disease. The cohort reflected a female to male ratio of 1:4. Approximately 47% of patients in this cohort received adequate BCG and risk stratification was performed as per the recent IBCG guidelines (**Supplementary data, Figure S3**). High-grade recurrence was observed in 33% patients in this cohort with 37 patients classified as BCG refractory and 124 as BCG responders. In this study, we used CD79a as the pan B cell marker. Kaplan-Meier survival analysis revealed that both female and male patients with high epithelial and stromal CD79a+ B cells in their pre-treatment tumors had significantly shorter RFS and PFS (**Figure 1A, 1B** and **S4**). Furthermore, comparison of the survival profiles specific to male and female patients revealed that females with high stromal density of CD79a+ B cells in their pre-BCG tumors experienced worse RFS and PFS (**Supplementary data, Figure S5**) compared to males. Quantification of immune cells in the stromal regions showed a significantly higher density of CD79a+ B cells (p=0.015) and CD163+ M2 macrophages (p=0.027) in BCG refractory patients compared to the responders (**Figure 1C and 1D**). Similar analysis of CD163+ cell densities within the epithelial compartment did not show any significant differences (**Supplementary data, Figure S6**).

**Figure 1.**
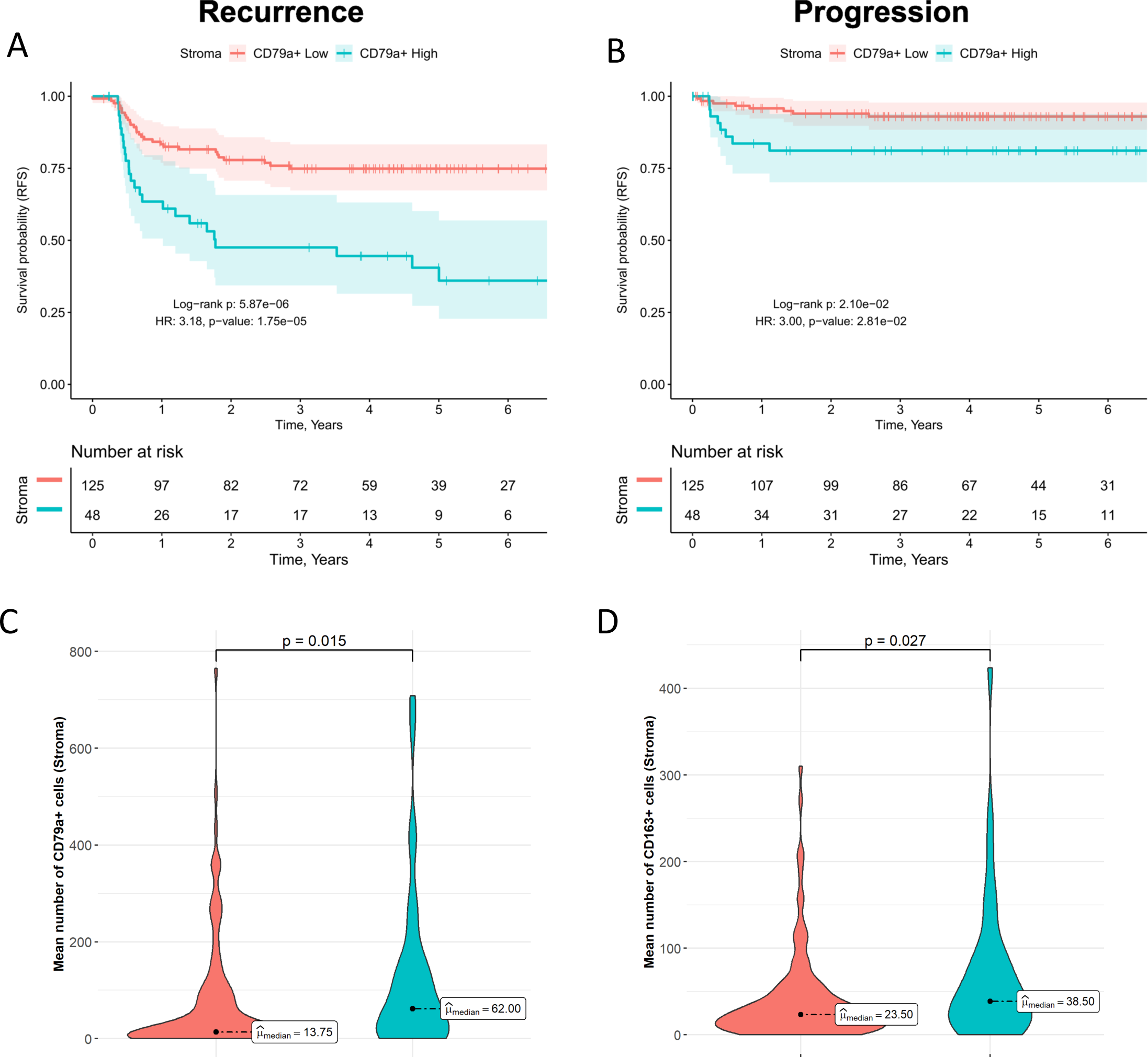
High intra-tumoral CD79a+ B cell density associates with early recurrence and progression post BCG immunotherapy. **A)** Patients with high stromal CD79a+ B cell density have significantly shorter RFS compared to patients with low density (Log-rank p=5.83e-6, IRR=0.29, HR=3.18, p=1.75e-5). **B)** Patients with high stromal CD79a+ B cell density have significantly shorter PFS compared to patients with low density (Log-rank p=2.07e-2, IRR=0.32, HR=3.00, p=2.8e-2), **C)** BCG refractory patients have significantly higher stromal CD79a+ cell density compared to BCG responders (W_Mann-Whitney_=1744.5, p=0.027), and **D)** BCG refractory patients have significantly higher stromal CD163 cell density compared to BCG responders (W_Mann-Whitney_=1688.00, p=0.015).

### Stromal CD79a+ B cell density is positively correlated with M2-like macrophages, CD8+ cytotoxic T cells, CD103+ cells and PD-1 immune checkpoint expression

We then evaluated the correlations between cell types and immune checkpoint expression in both the stromal and epithelial compartments. Spearman correlation analysis revealed significantly positive correlations between CD79a+ B cells and CD163+ M2-like macrophages (r=0.74, p=1×10^-7^), CD3+ CD8+ cytotoxic T cell (r=0.55, p=4×10^-4^), and PD-1 immune checkpoint expressing cells (r=0.55, p=4×10^-4^) in the stromal compartment of the BCG refractory group (**Figure 2A**). Similar correlations were observed in the BCG responder group in which CD79a+ B cells were significantly positively correlated with CD163+ M2 macrophages (r=0.61, p=3×10^-14^), CD3+CD8+ cytotoxic T cells (r=0.68, p=2×10^-10^), and both PD-1 (r=0.67, p=2×10^-16^) and PD-L1 (r=0.54, p=7×10^-11^) immune checkpoints (**Figure 2B**). Importantly, within the BCG refractory group, correlations between immune cell types in the epithelial compartments were not significant as observed within the stromal region (**Figures 2C and 2D**). In the stromal regions of tumors from BCG responder and refractory groups, CD163+ macrophages and CD8+ cytotoxic T cells also depicted moderate positive correlation with each other (r=0.67-0.68). Cytotoxic T cell density was highly correlated with the expression of PD-1 immune checkpoint (r=0.87-0.88) in both the groups.

**Figure 2.**
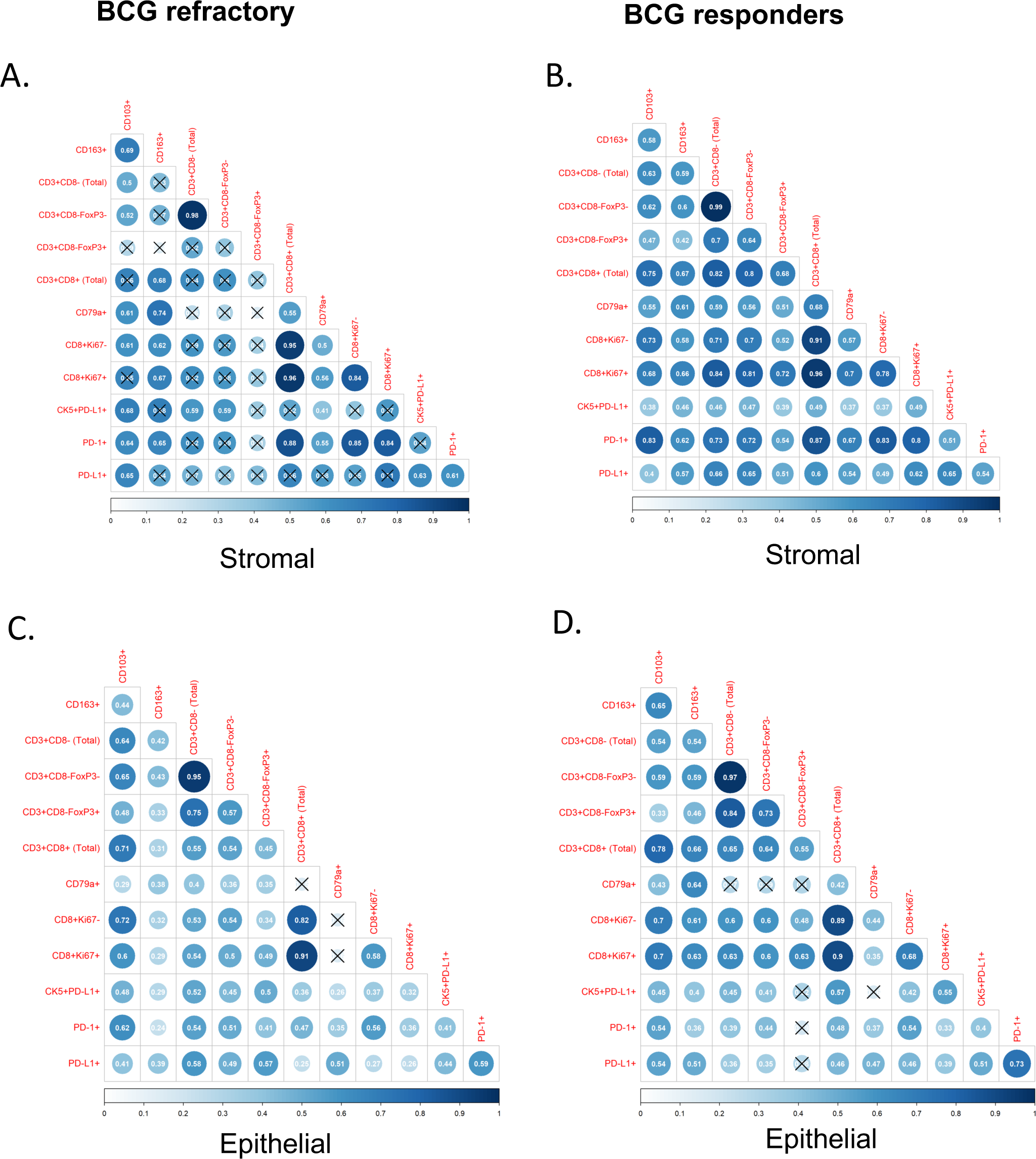
Stromal CD79a+ B cells are strongly correlated with CD163+ M2-like macrophages, CD8+ cytotoxic T cells, CD103+ cells and PD-1 immune checkpoint expression. Spearman correlation plot for immune cell populations in the stromal (A and B) and epithelial (C and D) compartments of tumors from BCG responders and refractory. Correlation plots showing the correlations between immune cells in tumor from patients categorized as BCG refractory and responders.

### Increased stromal density of CD8+ T cells, CD103+ cells and M2-like macrophages is associated with early recurrence post-BCG

Investigation of other immune cells in the stromal and epithelial compartment of each tumor core demonstrated a higher density of CD163+ M2-like macrophages, CD3+CD8+ cytotoxic T cells, and CD103+ cells in the BCG non-responder group. Kaplan-Meier survival analysis revealed a significant association between high stromal density of immune cells (M2-like macrophages, cytotoxic T cells and CD103+ cells) and overall shorter RFS (HR=2.87, p < 0.001, **Figure 3**).

**Figure 3.**
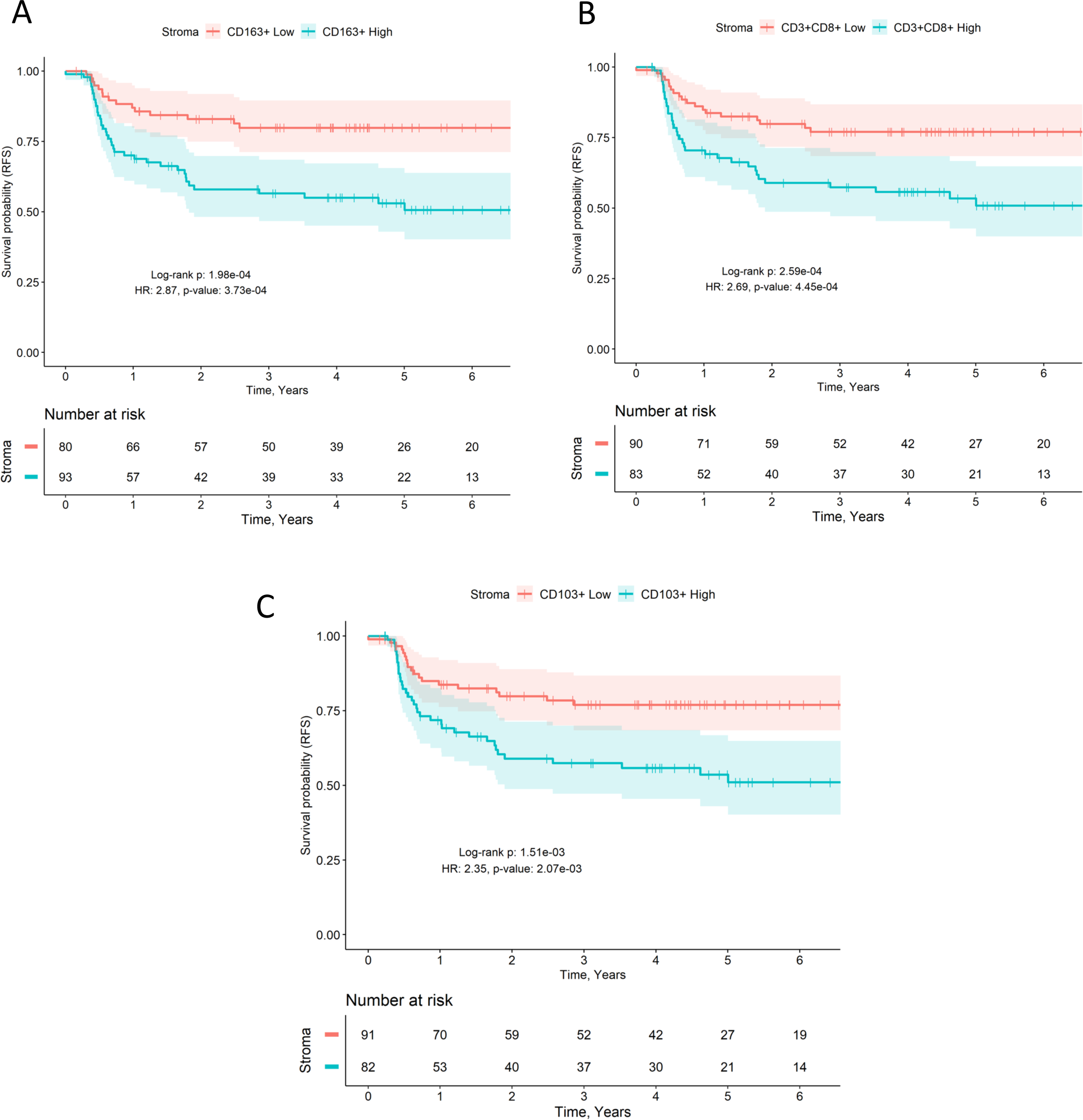
High tumor infiltrating M2-like macrophage, CD8+ and CD103+ T cell density associates with early recurrence post BCG immunotherapy. **A)** Patients with high stromal CD163+ macrophage density have significantly worse RFS compared to patients with low density. Log-rank p=1.98e-6, IRR=0.31, HR=2.87, p=3.73e-4. **B)** Patients with high stromal CD3+CD8+ cell density have significantly worse RFS compared to patients with low density. Log-rank p=2.59e-4, IRR=0.34, HR=2.69, p=4.45e-2. **C)** Patients with high stromal CD103+ cell density have significantly worse RFS compared to patients with low density. Log-rank p=1.51e-3, IRR=0.40, HR=2.35, p=2.07e-3

Spearman correlation analysis revealed that PD-1+ cell density is highly and significantly correlated with CD103+ (r=0.64-0.83), CD163+ M2-like macrophages (r=0.62-0.65), and both Ki67+ and Ki67-CD8+ T cells (r=0.8-0.85). Furthermore, significant correlations were observed between PD-1 and CD103+ cells (r=0.64-0.83), in the stromal compartment indicate heightened immune checkpoint expression in these cells. Similarly, a strong correlation (**Figure 2A and 2B**) was found between PD-1 expression and both proliferating (Ki67+) and non-proliferating (Ki67-) cytotoxic CD8 + T cells (r=0.8-0.85). High expression of PD-1 could potentially suggest T cell activation in BCG responders and a state of immune exhaustion in BCG refractory patients. These findings underscore the considerable variability in immune responses to BCG immunotherapy.

### Elevated density of stromal CD79a+ B cells, CD8+ T cells, CD103+ cells, and M2-like macrophages is associated with increased disease stage and heightened AUA risk score

To determine the stage and risk associated immune cell profiles, we performed a Pearson’s χ² analysis (**Supplementary data**, **Table S1**). Among the patients who received BCG treatment, a higher density of CD79a+ B cells, CD103+ cells, and M2-like macrophages in the stromal compartment of their tumors was associated with higher grade (χ²(1, N=173) = 10.23, p = 0.001, χ²(1, N=173) = 5.54, p = 0.019, χ²(1, N=173) = 5.95, p = 0.015, respectively) and stage (χ²(1, N=173) = 6.11, p = 0.013, χ²(1, N=173) = 9.55, p = 0.002, respectively), except for the higher CD79+ B cell density. The basal subtype was notably more prevalent in patients with an elevated stromal density of CD79a+ B cells and CD103+ cells, as confirmed by a Fisher’s Exact test (p = 0.016 and 0.025, respectively).

### Univariable and multivariable analysis reveals high pre-BCG tumor immune cell density is associated with a higher risk of recurrence and progression

A CPH based analysis revealed significant associations between immune cell infiltration profiles and clinical outcomes (**Figure 4**). Specifically, in the epithelium PD-1+ (HR_SD_=1.223, p=0.026), CD79a+ B cells (HR_SD_=1.198, p=0.024), and CK5+ PD-L1+ cancer (HR_SD_=1.241, p=0.019) cell density was associated with worse RFS (**Supplementary data Table S2**). CD79a+ B cells (HR_SD_=1.324, p=0.022), CD163+ M2-like macrophages (HR_SD_=1.578, p < 0.001), CK5+PD-L1+ cancer cells (HR_SD_=1.391, p=0.01), and PD-1+ immune checkpoint expressing cells (HR_SD_=1.352, p=0.017), in the epithelial compartment, were also associated with worse PFS (**Supplementary data Table S3**).

**Figure 4.**
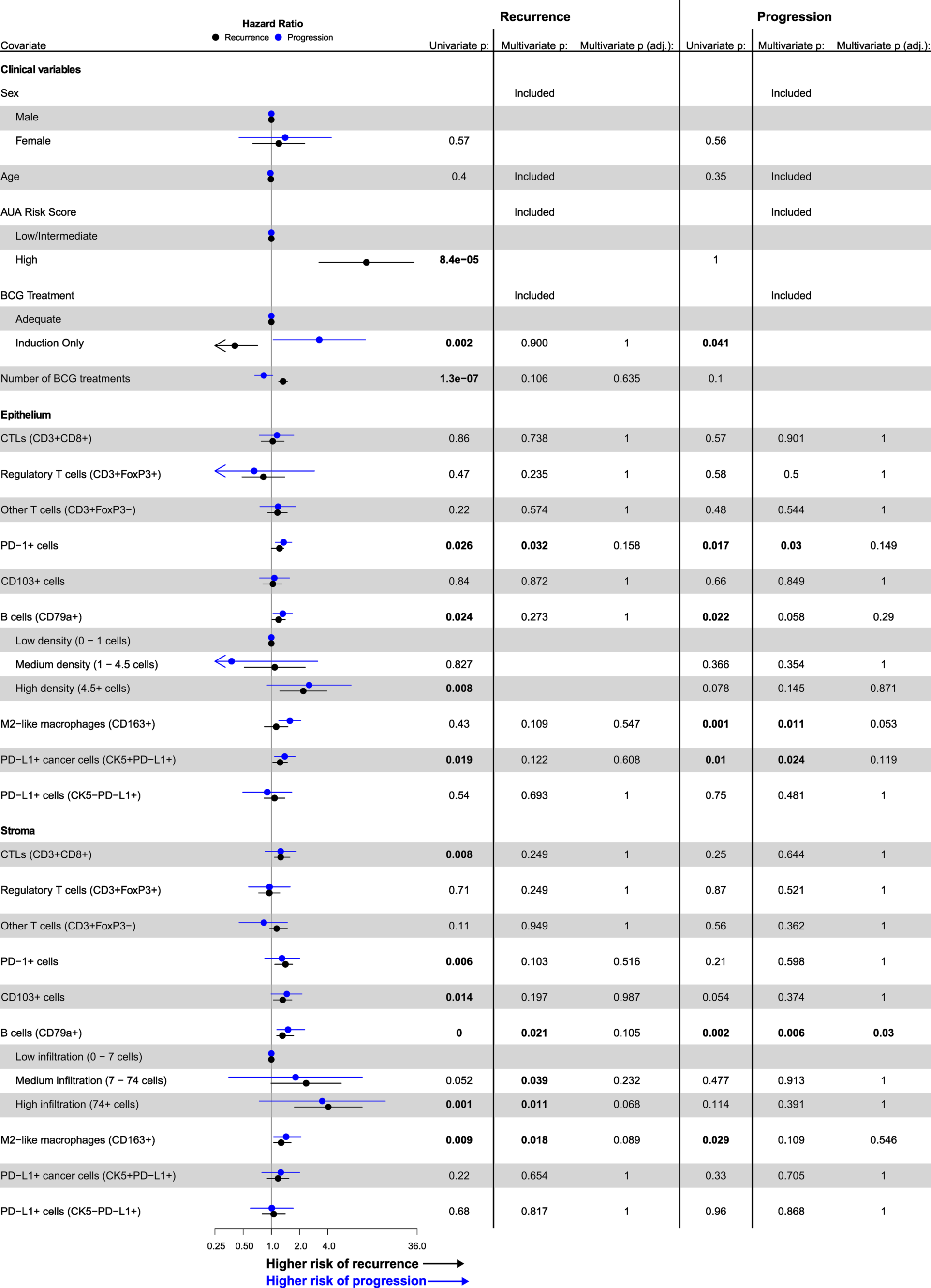
Forest plots demonstrating patient demographics and immune markers with clinical outcomes post BCG immunotherapy. Forest plot based showing univariate Cox Proportional Hazards (CPH) regression analyses, with recurrence highlighted in black and progression in blue. Dots represent hazard ratios and lines represent the 95% confidence intervals. Hazard ratios for immune cell types are displayed graphically in terms of risk per standard deviation (σ) increment in cell infiltration. Multivariable CPH regression analyses were performed for each immune phenotype separately, including sex, age, AUA risk score, and amount of BCG treatment received after the analyzed tumor. P values were calculated using the Wald test, and multivariable p values were adjusted using the Bonferroni correction.

In the stroma, CD3+CD8+ cytotoxic T cells (HR_SD_=1.258, p=0.008), CD79a+ B cells (HR_SD_=1.315, p < 0.001), CD163+ M2-like macrophages (HR_SD_=1.274, p=0.009), CD103+ cells (HR_SD_=1.322, p=0.014), and PD-1+ immune checkpoint expressing cells (HR_SD_=1.415, p=0.006) were associated with worse RFS. Only CD79a+ B cells (HR_SD_=1.508, p=0.002), and CD163+ M2-like macrophages (HR_SD_=1.438, p=0.029), in the stromal compartment, were significantly associated with PFS.

In a multivariable model considering sex, age, AUA risk score, and BCG treatment adequacy as covariates, none of the immune cells in either the stromal or epithelial compartment were significantly associated with a change in RFS after adjustment for multiple hypothesis testing. CD79a+ B cells, in the stromal compartment, were however significantly associated with worse PFS (HR_SD_=1.663, p=0.03), with and without Bonferroni correction.

## Discussion

Recurrence or progression to muscle invasive disease following BCG immunotherapy remain the major challenges in the management of patients with NMIBC. To determine the factors that lead to a poor response, studies have extensively examined the treatment-naïve TIME in tumors collected at TURBT from patients with NMIBC. Specifically, these prior investigations have focused on T cells and myeloid cells and have significantly advanced the current state of knowledge on the complex nature of the TIME in NMIBC tumors [19, 20, 23, 24]. However, clinical translation to actionable biomarkers is still unavailable. Additionally, B cells that constitute the key cellular mediators of mucosal immunity, via their dual functions as antigen presenting and antibody producing cells, remain understudied in the context of response to BCG immunotherapy.

A significant and novel observation from the current study is the association between a high density of stromal and intra-epithelial B cells in the BCG naïve tumors from patients who experienced early recurrence or progression to muscle invasive disease. Our findings demonstrate a robust association between CD79a+ B cell infiltration in both stromal and epithelial compartments, which is more pronounced in high-grade tumors from female patients who experienced early recurrence and disease progression following BCG therapy. These observations align with current literature highlighting the importance of CD79a+ B cells in driving tumor progression and fostering resistance to BCG therapy, particularly within the stromal milieu [21].

A higher stromal B cell density and a positive correlation between densities of CD79a+ B cells, M2-like macrophages, CD103+ cells, PD-1 immune checkpoint expression, compared to that of the epithelial compartment, is indicative of chronic inflammation and potential exhaustion within or outside of either stromal immune cell aggregates or bladder mucosa associated lymphoid tissue referred to as tertiary lymphoid structures (TLSs). Mucosa associated TLSs evolve as a result of chronic inflammation such as carcinogen exposure or persistent antigenic stimuli including recurrent urinary tract infection or autoimmune mediated Hunner’s lesion interstitial cystitis [25]. Supporting this, our previous report demonstrated the compounding effects of aging and chronic carcinogenesis associated inflammation on increased B cell recruitment and TLS formation in the urinary bladder [26, 27]. We also showed the higher expression of multiple immune exhaustion associated proteins within TLSs and their peri-tumoral location associated with higher stage and poor response to BCG [26, 28]. A bladder tumor associated chronic inflammatory state also suggests an exhausted pre-BCG host immune system. Consequent to such systemic immune exhaustion, exposure to BCG bacteria at weekly intervals during the induction phase of treatment potentially intensifies the chronic systemic inflammatory state, specifically in patients with higher stromal B cell profiles in their pre-BCG tumors. This is supported by a recent study conducted by Strangaard *et al.,* [20] which examined pre- and post-BCG paired tumor samples and showed T cell exhaustion as the factor underlying a BCG unresponsive disease. Overall, these findings suggest that in a subset of patients who do not benefit from BCG, signatures of innate and adaptive immune exhaustion within the pre-BCG tumor immune profiles could inform patient selection for immunomodulatory BCG treatment.

Most recently, de Jong *et al*., reported the presence of three BCG response subtypes (BRS1, BRS2, and BRS3) of pre-treatment tumors that show a distinct association with progression of NMIBC to muscle invasive disease [21]. Specifically, in this study based on bulk RNA-sequencing of tumors, the BRS3 subtype enriched with immune gene signatures was shown to associate with poor outcomes following BCG treatment. Tumors with a BRS3 subtype (increased expression of genes associated with B cell, M1/M2 macrophage, CD8+ T cell, T_reg_ function and immune exhaustion) exhibited the most aggressive behavior and early progression. Notably, luminal subtype tumors that are known to exhibit a low immune infiltration profile, were not enriched for a BRS3 gene signature. Unsurprisingly, the post-BCG recurrent tumors also carried the BRS3 phenotype and a basal tumor subtype [21]. In the current study, we used the pan B cell marker - CD79a, to identify tumor infiltrating B cells. Evaluation of bulk RNA-Seq profiles from the publicly available tumor transcriptomic profiles of 283 patients generated by de Jong *et al.,* validated our findings on the increased expression of *CD79A* and *CD163* transcripts in tumors from patients who experienced early progression to muscle invasive disease following BCG therapy.

Our counterintuitive findings showing inverse correlations between higher density of cytotoxic T cells and poor response to BCG immunotherapy, are also in alignment with previous literature suggesting the development of adaptive immune resistance following treatment in BCG non-responders [19, 21]. Previous studies have highlighted the involvement of regulatory B cell subsets in fostering an immunosuppressive environment conducive to immune evasion and tumor progression [29–31]. Remarkably, our study corroborates these prior findings, establishing a positive correlation between CD79a+ B cells and CD163+ M2-like macrophages, CD8+ Ki67+ proliferating cytotoxic T-cells, and PD-1 expression within the stromal compartment of BCG-refractory patients.

The notable correlation between CD79a+ B-cells, CD8+ T cells and the PD-1 immune checkpoint protein suggests a potential regulatory role exerted by immune checkpoint mechanisms in modulating the functional activity of cytotoxic T cells in response to BCG therapy. Similarly, the presence of CD8+ T cells and CD103+ cells implies an actively mounting immune response against the tumor. However, their correlation with shorter RFS strongly suggests the possibility of dysfunction or exhaustion, potentially arising from chronic antigen exposure or inhibitory signals from other immune cell subsets. Immune cells expressing the α_E_ integrin CD103 (mainly T_RM_ cells) are widely distributed across mucosal surfaces [32]. In this study we did not use a T cell phenotypic marker to define CD103+ cell, however, their significantly high correlation with CD8+ cells and previous reports confirming the T cell specific intra-tumoral expression pattern of CD103 and the chronic nature of the disease, is suggestive of their T_RM_ phenotype [33, 34]. Indeed, the expression of CD103 has been reported on multiple cell types (Tregs, tissue resident memory T cells) within the stromal compartment across various cancer types [35]. Future studies should further characterize their effector memory and central memory functional states.

Overall, these findings from multiple independent cohorts confirm that spatial organization and density of B cells within the pre-BCG TIME could be developed into a robust clinically feasible assay for application on TURBT specimens prior to initiation of BCG treatment.

An important and novel finding from our study is the significantly worse RFS and PFS in BCG treated female patients with high stromal and intra-epithelial B cell density compared to that of males. As reported by us and others, the physiologically higher B cell frequency in females compared to that of males is likely a key factor underlying the higher degree of their trafficking to the bladder TIME upon chronic antigenic stimulation. Previously, we have reported a significantly shorter PFS in females based on evaluation of two large independent cohorts of patients with NMIBC. We also showed that irrespective of the treatment type, a higher pre-treatment intra-tumoral B cell density is associated with early RFS and PFS in patients with NMIBC [18].

While these findings indicate that B cell proportions increase with disease progression and due to the immune stimulatory effects of BCG bacteria, we speculate that upon treatment with cytotoxic chemotherapy such as gemcitabine-docetaxel chemotherapy or immune checkpoint inhibition, such responses may lead to a favorable response via immunogenic cell death mediated events. Such associations have been reported in other cancers and in muscle invasive bladder cancer, where higher pre-treatment B cell density associated with positive outcomes following treatment [36–38]. Finally, in this study, we observed a significant association between increased density of CDK5+ PDL-1+ cells indicative of an aggressive cancer cell phenotype and shorter PFS. Constitutive expression of PD-L1 on cancer cells could result from mutations in DNA damage repair genes or could be induced via the phenomenon of adaptive immune resistance during disease progression [39]. While the mutational profiles of tumors used in our study were not available, previous studies have reported an enrichment of DNA damage repair gene mutations in high-grade NMIBC tumors [40, 41].

This study is not without limitations. We acknowledge that tumor cores of 0.6 - 1 mm diameter in a TMA do not capture the heterogeneity within the larger tumor landscape. Recently, Lindskrog *et al*., [23] reported the presence of multiple tumor subtypes within a single tumor specimen. This finding supports a potential caveat of TMA based studies and the need for surveying whole tumor sections to derive a comprehensive spatial organization. However, with regard to our previous report confirming stromal B cell infiltration profiles, and TLSs in whole tumor sections of BCG non-responders, we posit that a careful selection of tissue cores capturing tumor invasive margins is required for larger cohort-based biomarker studies. A deeper characterization of the B cell subsets and their cellular neighborhoods using whole tumor sections is also needed to gain further insights into their antigen presenting and/or antibody producing functions. Finally, exploration of the finer spatial relationships between the several immune cell populations and tumor invasive margins within pre- and post-BCG treatment tumors is critical to determine if the localization of specific cellular phenotypes associate with BCG outcome.

To conclude, the findings presented in this study advance the current understanding of the immune landscape characterizing the NMIBC microenvironment and its association with patient responses to BCG immunotherapy. Ultimately, these findings have implications not only for the development of novel therapeutic strategies targeting B cell specific exhaustion but also to inform approaches for precise sequencing of drugs targeting the immune checkpoint pathways to improve the efficacy of treatments for NMIBC. The results also contribute to a growing body of literature on sex differences in the immune cell landscape and the importance of sex considerations in BCG immunotherapy and the need for a sex-disaggregated data analyses approach in biomarker studies to address the interpretations associated with inconsistent findings across different cohorts. Given the advent of newer immunotherapies as well as gemcitabine-docetaxel chemotherapy in the BCG naïve setting, early identification of patients who may not benefit from BCG, using information from pre-treatment immune landscape, will allow precise incorporation of newer therapeutics in NMIBC.

## Author contributions and acknowledgements

MK conceptualized the overall study and contributed to study design, manuscript writing and revision. DRS and BR contributed to the study design. SC performed manual validation of mIF staining and automated scoring. BR performed all data analysis relating to the KHSC cohort and contributed to manuscript writing and revision. PY, SR and KP contributed to manuscript writing and revision. KS, HG and AG performed additional analysis included in this study, such as analysis for the independent validation cohort and subtyping and contributed to manuscript writing and reviewing. CJ and DMB provided the KHSC tissue cohort and associated clinical data. KT guided all data analysis. DRS and CJ helped with clinical classifications of BCG treated patient specimens. All authors reviewed the manuscript. PY is supported by Robert Wilson Fellowship for PhD training. This study was supported by research operating grants from Bladder Cancer Canada (MK, DRS and DMB), Cancer Research Society (MK, DRS and DMB), Ontario Institute for Cancer Research/Ontario Molecular Pathology Research Network (DMB); Ontario Ministry of Research Innovation and Science: Early Researcher Award and Canada Foundation for Innovation (MK). We thank Shakeel Virk at the QLMP for his assistance with imaging and Halo software and Katy Milne at BC Cancer Agency’s Molecular and Cellular Immunology Core for immunofluorescence staining of TMAs.

