## Supplementary data for "Increased stromal densities of B cells, CD103+ cells, and CD163+ M2-like macrophages associate with poor clinical outcomes in BCG treated non-muscle invasive bladder cancer"

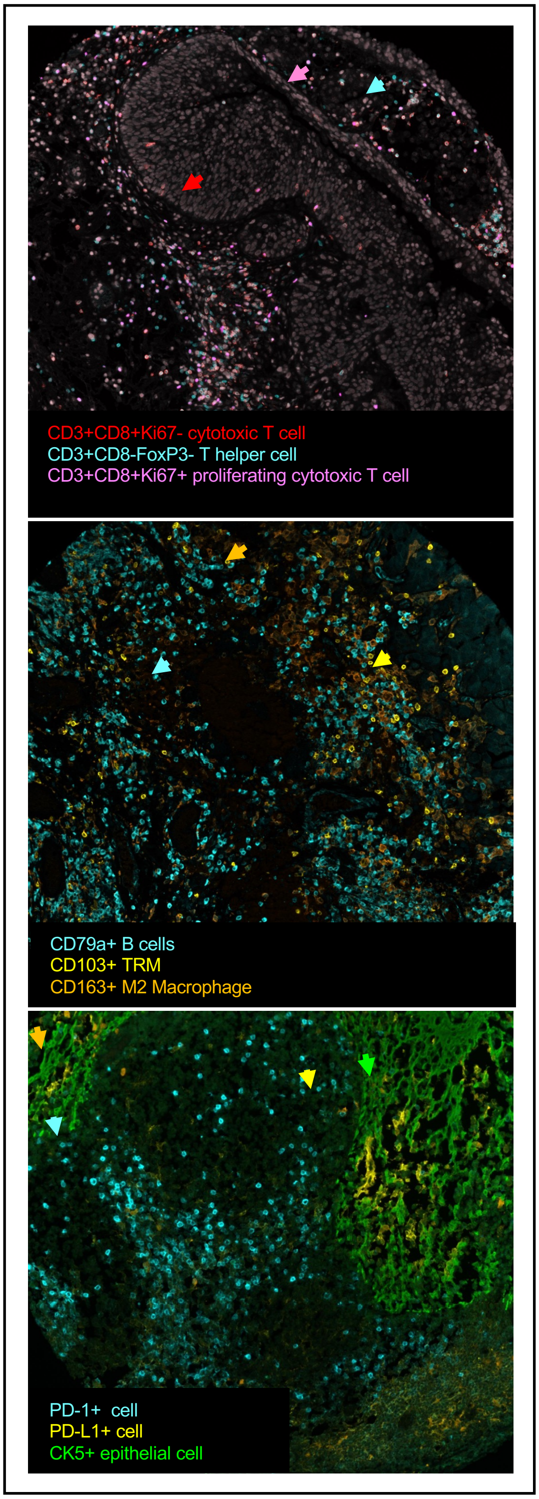

**Figure S1**. Representative images of multiplex immunofluorescence stained tissue microarray cores showing the cells of interest across the three panels of antibodies.

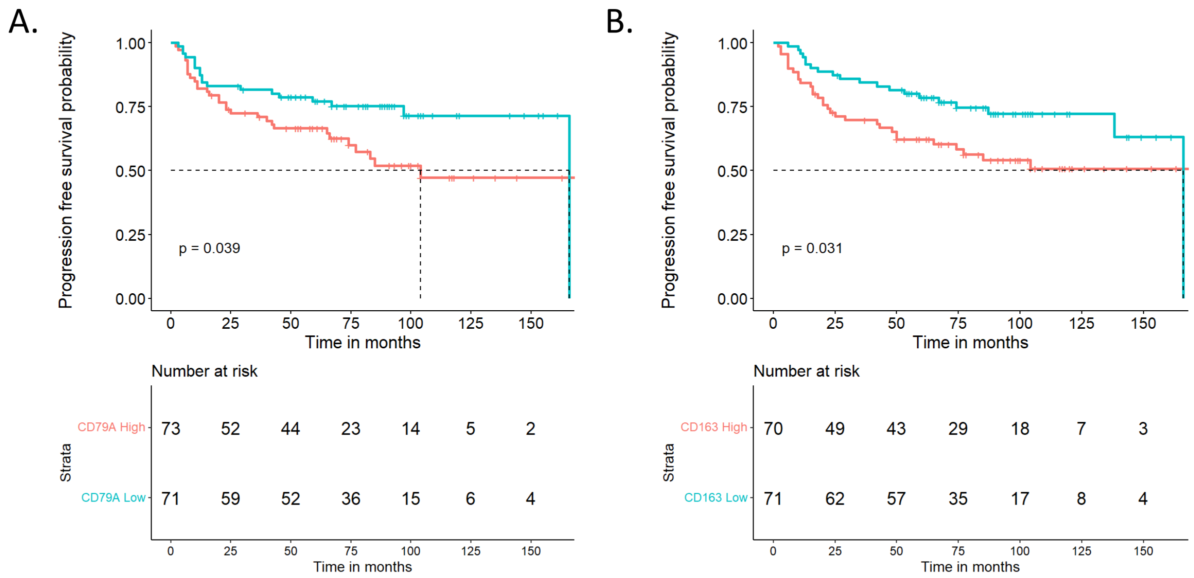

**Figure S2. Increased expression of *CD79A* and *CD163* transcripts in high-grade pre-BCG tumors associates with shorter progression free survival in patients treated with BCG.** Kaplan-Meier survival curve showing significantly shorter PFS in patients (independent cohort, n=283) with higher expression of *CD79A* **(A)** and *CD163* **(B)** transcripts in their pre-BCG tumors.

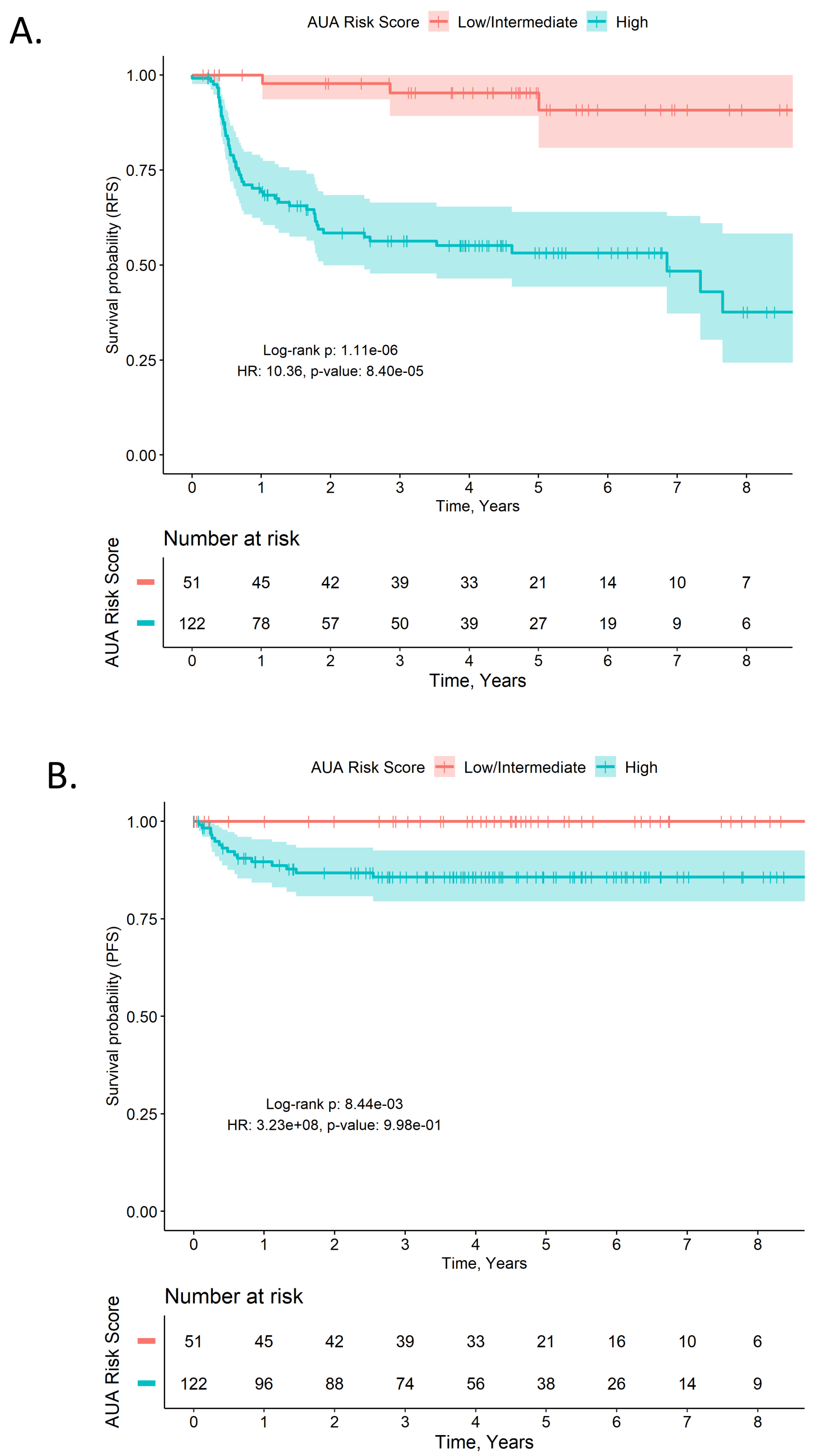

**Figure S3**. **A)** Patients classified as high risk according to the AUA risk stratification guidelines have a significantly worse RFS compared to patients classified as low or intermediate risk. Log-rank test p=1.11e-6, IRR=0.08, HR=10.36, p=8.40e-5. **B)** Patients classified as high risk according to the AUA risk stratification guidelines have a significantly worse PFS compared to patients classified as low or intermediate risk. Log-rank test p=8.44e-3, IRR=0, HR=3.23e8, p=1

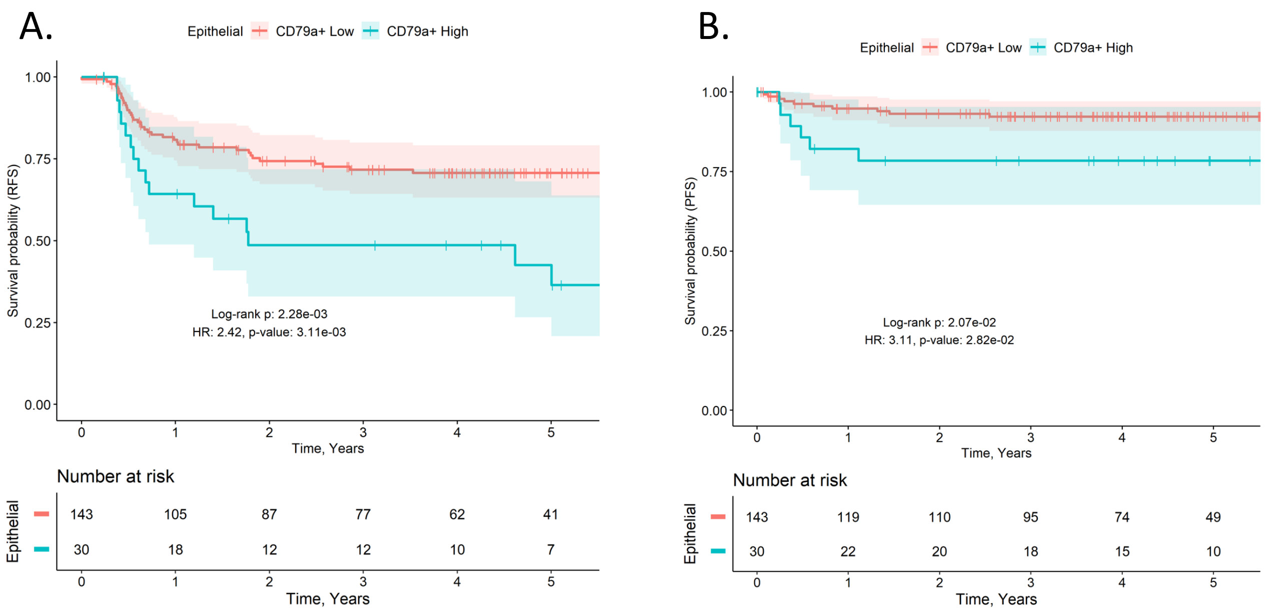

**Figure S4. Increased epithelial CD79a cell density associates with shorter recurrence and progression free survival. A)** Patients with high epithelial CD79a+ cell density have significantly worse RFS compared to patients with low density. Log-rank p=2.28e-3, IRR=0.37, HR=2.42, p=3.11e-3. **B)** Patients with high epithelial CD79a+ density have significantly worse PFS compared to patients with low density. Log-rank p=2.07e-2, IRR=0.30, HR=3.11, p=2.82e-2

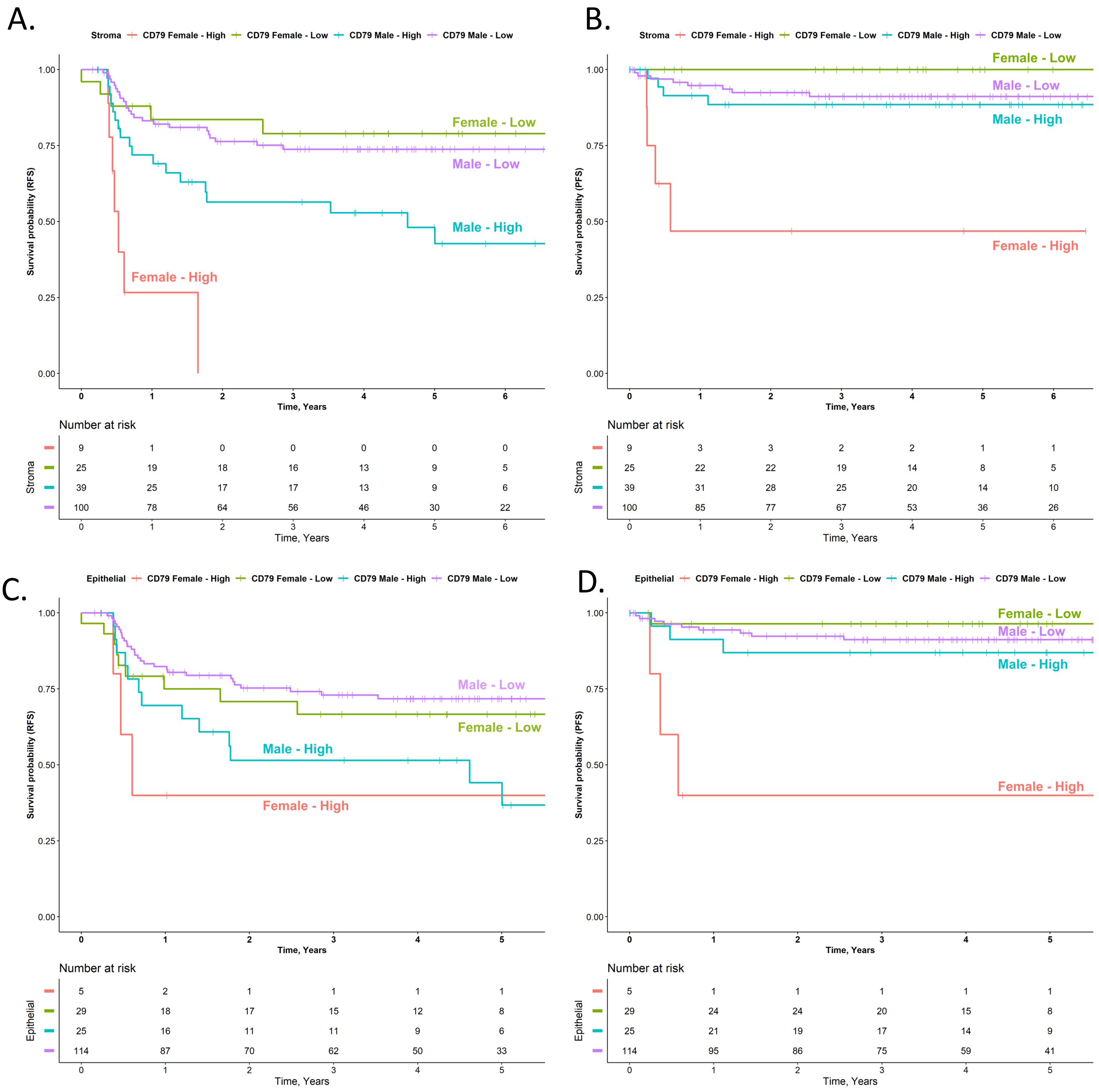

**Figure S5. Sex associated profiles of stromal and epithelial CD79a+ B cell density and clinical outcomes post BCG therapy. A)** Females with high stromal CD79a+ B cell density have significantly lower RFS compared to females with low density (log-rank p=3.90e-4; HR=8.87, p=1.07e-3). Males with high stromal CD79a+ B cell density have significantly lower RFS compared to males with low density (log-rank p=3.20e-3; HR=2.48, p=2.93e-3). **B)** Females with high stromal CD79a+ B cell density have significantly lower PFS compared to females with low density (log-rank p=1.70e-4; HR=1.08e10, p=0.999). **C)** Females with high epithelial CD79a+ density have significantly lower RFS compared to females with low density (log-rank p=0.041; HR=2.40, p=8.78e-3). **D)** Females with high epithelial CD79a+ B cell density have significantly lower PFS compared to females with low density (log-rank p=5.90e-4; HR=20.66, p=8.85e-3).

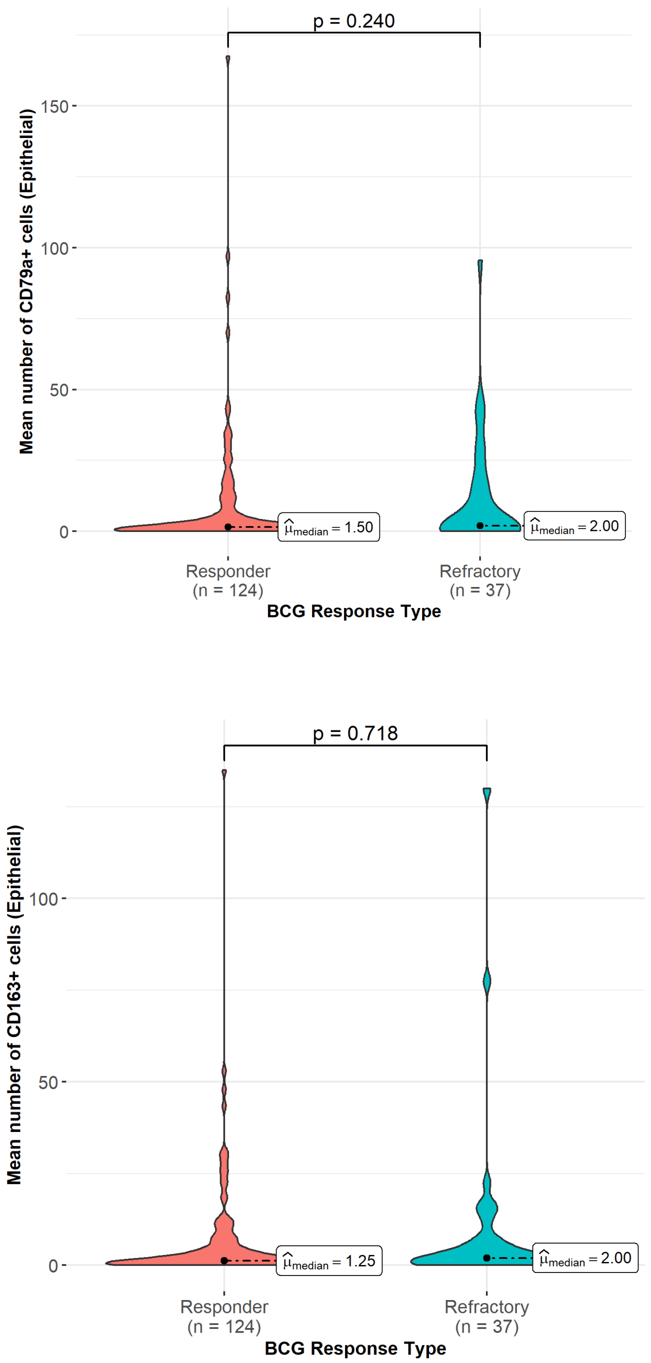

**Figure S6. Distribution of CD79a+ B cells and CD163+ M2-like macrophages within epithelial regions of BCG responders and refractory patients.**

**Supplementary Table S1. Stage and grade specific profiles of CD79a+ B cell density in pre-treatment tumors from female and male patients**

A

| | Male | Female | Overall |
|---|---|---|---|
| | (N=139) | (N=34) | (N=173) |
| **Initial Stage/Grade** | | | |
| **CIS** | 11 (7.9%) | 2 (5.9%) | 13 (7.5%) |
| **CD79a+ cells** | 131 | 15.5 | |
| **PUNLMP** | 1 (0.7%) | 0 (0%) | 1 (0.6%) |
| **CD79a+ cells** | 2 | - | |
| **T1HG** | 37 (26.6%) | 14 (41.2%) | 51 (29.5%) |
| **CD79a+ cells** | 147.1 | 178.7 | |
| **TaHG** | 63 (45.3%) | 10 (29.4%) | 73 (42.2%) |
| **CD79a+ cells** | 61.1 | 40.8 | |
| **TaLG** | 25 (18.0%) | 8 (23.5%) | 33 (19.1%) |
| **CD79a+ cells** | 27.5 | 12.5 | |
| **Unknown** | 2 (1.4%) | 0 (0%) | 2 (1.2%) |
| **CD79a+ cells** | 41.5 | - | |
| **Stromal CD79a+ cell density** | | | |
| **Mean** | 82.8 | 89.4 | 84.1 |
| **Stromal CD79a+ cell density (Categorized)** | | | |
| Low (0-7) | 49 (35.3%) | 10 (29.4%) | 59 (34.1%) |
| Medium (7-74) | 45 (32.4%) | 13 (38.2%) | 58 (33.5%) |
| High (74-765) | 45 (32.4%) | 11 (32.4%) | 56 (32.4%) |
| **Stromal CD103+ cells** | | | |
| **Mean** | 19.6 | 22.2 | 20.1 |
| **Stromal CD163+ cells** | | | |
| **Mean** | 42.7 | 58.4 | 45.8 |

B

| Stromal B cell density | T1HG | TaHG | TaLG | CIS |
|---|---|---|---|---|
| **High (n=57)** | 28 (49%) | 18 | 5 | 5 |
| **Med (n=58)** | 16 (27%) | 26 | 7 | 7 |
| **Low (n=59)** | 7 (11%) | 29 | 21 | 1 |

| | Univariate | | | Multivariate | | |
|---|---|---|---|---|---|---|
| Variable | HR | HR (SD) | P-Value | HR | P-Value | Adj. P-Value |
| **Clinical variables** | | | | | | |
| **Sex** | | | | | Included | |
| **Male** | 1.000 | 1.000 | | | | |
| **Female** | 1.202 | 1.000 | 0.570 | | | |
| **Age** | 0.989 | 1.000 | 0.400 | | Included | |
| **AUA Risk Score** | | | | | Included | |
| **Low/Intermediate** | 1.000 | 1.000 | | | | |
| **High** | 10.360 | 1.000 | 0.000 | | | |
| **BCG Treatment** | | | | | Included | |
| **Adequate** | 1.000 | 1.000 | | | | |
| **Induction Only** | 0.408 | 1.000 | 0.002 | 0.953 | 0.900 | 1.000 |
| **Number of BCG treatments** | 1.335 | 1.000 | 0.000 | 1.621 | 0.106 | 0.635 |
| **Epithelium** | | | | | | |
| **CTLs (CD3+CD8+)** | 1.001 | 1.035 | 0.860 | 1.001 | 0.738 | 1.000 |
| **Regulatory T cells (CD3+FoxP3+)** | 0.986 | 0.824 | 0.470 | 0.973 | 0.235 | 1.000 |
| **Other T cells (CD3+FoxP3-)** | 1.010 | 1.163 | 0.220 | 1.005 | 0.574 | 1.000 |
| **PD-1+ cells** | 1.002 | 1.223 | 0.026 | 1.002 | 0.032 | 0.158 |
| **CD103+ cells** | 1.001 | 1.038 | 0.840 | 1.001 | 0.872 | 1.000 |
| **B cells (CD79a+)** | 1.009 | 1.198 | 0.024 | 1.005 | 0.273 | 1.000 |
| **Low density (0 - 1 cells)** | 1.000 | 1.000 | | | | Reference |
| **Medium density (1 - 4.5 cells)** | 1.087 | 1.000 | 0.827 | | | |
| **High density (4.5+ cells)** | 2.195 | 1.000 | **0.008** | | | |
| **M2-like macrophages (CD163+)** | 1.007 | 1.127 | 0.430 | 1.013 | 0.109 | 0.547 |
| **PD-L1+ cancer cells (CK5+PD-L1+)** | 1.044 | 1.241 | **0.019** | 1.033 | 0.122 | 0.608 |
| **PD-L1+ cells (CK5-PD-L1+)** | 1.018 | 1.082 | 0.540 | 0.988 | 0.693 | 1.000 |
| **Stroma** | | | | | | |
| **CTLs (CD3+CD8+)** | 1.003 | 1.258 | **0.008** | 1.002 | 0.249 | 1.000 |
| **Regulatory T cells (CD3+FoxP3+)** | 0.998 | 0.952 | 0.710 | 0.993 | 0.249 | 1.000 |
| **Other T cells (CD3+FoxP3-)** | 1.001 | 1.141 | 0.110 | 1.000 | 0.949 | 1.000 |
| **PD-1+ cells** | 1.004 | 1.415 | **0.006** | 1.002 | 0.103 | 0.516 |
| **CD103+ cells** | 1.011 | 1.322 | **0.014** | 1.007 | 0.197 | 0.987 |
| **B cells (CD79a+)** | 1.002 | 1.315 | **0.000** | 1.002 | 0.021 | 0.105 |
| **Low infiltration (0 - 7 cells)** | 1.000 | 1.000 | | | | Reference |
| **Medium infiltration (7 - 74 cells)** | 2.351 | 1.000 | 0.052 | 2.523 | 0.039 | 0.232 |
| **High infiltration (74+ cells)** | 4.059 | 1.000 | **0.001** | 2.984 | 0.011 | 0.068 |
| **M2-like macrophages (CD163+)** | 1.004 | 1.274 | **0.009** | 1.004 | 0.018 | 0.089 |
| **PD-L1+ cancer cells (CK5+PD-L1+)** | 1.033 | 1.179 | 0.220 | 1.014 | 0.654 | 1.000 |
| **PD-L1+ cells (CK5-PD-L1+)** | 1.011 | 1.059 | 0.680 | 0.993 | 0.817 | 1.000 |

**Supplementary Table S3:** Cox Proportional Hazards analysis results for recurrence free survival.

| | Univariate | | | Multivariate | | | |
|---|---|---|---|---|---|---|---|
| Variable | HR | HR (SD) | P-Value | HR | HR (SD) | P-Value | Adj. P-Value |
| **Clinical variables** | | | | | | | |
| **Sex** | | | | | | Included | |
| **Male** | 1.000 | 1.000 | Reference | | | | |
| **Female** | 1.405 | 1.000 | 0.560 | | | | |
| **Age** | 0.978 | 1.000 | 0.350 | | | Included | |
| **AUA Risk Score** | | | | | | Included | |
| **Low/Intermediate** | 1.000 | 1.000 | Reference | | | | |
| **High** | 3.227E+08 | 1.000 | 1 | | | | |
| **BCG Treatment** | | | | | | Included | |
| **Adequate** | 1.000 | 1.000 | Reference | | | | |
| **Induction Only** | 3.248 | 1.000 | 0.041 | | | | |
| **Number of BCG treatments** | 0.831 | 1.000 | 0.1 | | | | |
| **Epithelium** | | | | | | | |
| **CTLs (CD3+CD8+)** | 1.004 | 1.149 | 0.570 | 1.001 | 1.026 | 0.901 | 1.000 |
| **Regulatory T cells (CD3+FoxP3+)** | 0.970 | 0.656 | 0.580 | 0.966 | 0.625 | 0.500 | 1.000 |
| **Other T cells (CD3+FoxP3-)** | 1.011 | 1.181 | 0.480 | 1.009 | 1.139 | 0.544 | 1.000 |
| **PD-1+ cells** | 1.003 | 1.352 | **0.017** | 1.003 | 1.316 | 0.030 | 0.149 |
| **CD103+ cells** | 1.002 | 1.078 | 0.660 | 0.999 | 0.961 | 0.849 | 1.000 |
| **B cells (CD79a+)** | 1.014 | 1.324 | **0.022** | 1.013 | 1.308 | 0.058 | 0.290 |
| **Low density (0 - 1 cells)** | 1.000 | 1.000 | | | | Reference | |
| **Medium density (1 - 4.5 cells)** | 0.377 | 1.000 | 0.366 | 0.364 | 1.000 | 0.354 | 1.000 |
| **High density (4.5+ cells)** | 2.533 | 1.000 | 0.078 | 2.252 | 1.000 | 0.145 | 0.871 |
| **M2-like macrophages (CD163+)** | 1.027 | 1.578 | **0.001** | 1.024 | 1.510 | 0.011 | 0.053 |
| **PD-L1+ cancer cells (CK5+PD-L1+)** | 1.068 | 1.391 | **0.010** | 1.064 | 1.364 | 0.024 | 0.119 |
| **PD-L1+ cells (CK5-PD-L1+)** | 0.978 | 0.905 | 0.750 | 0.950 | 0.795 | 0.481 | 1.000 |
| **Stroma** | | | | | | | |
| **CTLs (CD3+CD8+)** | 1.003 | 1.258 | 0.250 | 1.001 | 1.094 | 0.644 | 1.000 |
| **Regulatory T cells (CD3+FoxP3+)** | 0.998 | 0.957 | 0.870 | 0.993 | 0.844 | 0.521 | 1.000 |
| **Other T cells (CD3+FoxP3-)** | 0.999 | 0.831 | 0.560 | 0.998 | 0.735 | 0.362 | 1.000 |
| **PD-1+ cells** | 1.003 | 1.298 | 0.210 | 1.001 | 1.129 | 0.598 | 1.000 |
| **CD103+ cells** | 1.015 | 1.462 | 0.054 | 1.007 | 1.186 | 0.374 | 1.000 |
| **B cells (CD79a+)** | 1.003 | 1.508 | **0.002** | 1.004 | 1.663 | **0.006** | **0.030** |
| **Low infiltration (0 - 7 cells)** | 1.000 | 1.000 | | | | Reference | |
| **Medium infiltration (7 - 74 cells)** | 1.813 | 1.000 | 0.477 | 1.097 | 1.000 | 0.913 | 1.000 |
| **High infiltration (74+ cells)** | 3.495 | 1.000 | 0.114 | 1.979 | 1.000 | 0.391 | 1.000 |
| **M2-like macrophages (CD163+)** | 1.01 | 1.438 | **0.029** | 1.005 | 1.333 | 0.109 | 0.546 |
| **PD-L1+ cancer cells (CK5+PD-L1+)** | 1.05 | 1.262 | 0.330 | 1.017 | 1.090 | 0.705 | 1.000 |
| **PD-L1+ cells (CK5-PD-L1+)** | 1.00 | 1.011 | 0.960 | 0.991 | 0.953 | 0.868 | 1.000 |

**Supplementary Table S2:** Cox Proportional Hazards analysis results for progression free survival.
